# GLP-1 receptor agonist exenatide uncouples food intake from hedonic and anticipatory regulation in non-human primates: Insights from an operant meal schedule paradigm

**DOI:** 10.1101/2024.03.18.585513

**Authors:** Balázs Knakker, Judit Inkeller, Péter Kovács, Balázs Lendvai, István Hernádi

## Abstract

Glucagon-like peptide 1 (GLP-1), a neuroendocrine signal of energy balance and satiety, has a major role in regulating food intake behaviour. Here we investigated the effects of the GLP-1 agonist exenatide on palatability-driven feeding regulation in adult male rhesus macaques (n=5) using a novel operant food intake paradigm with four meal schedule conditions where two types of pellets with different palatability values were offered as meal in all combinations in two consecutive daily feeding sessions (S1 and S2). In control conditions, a strong, palatability-driven anticipatory effect was found in S1, followed by a complementary positive contrast effect in S2. After acute subcutaneous treatment with 1 µg/kg dose of exenatide 1 h before S1, food intake decreased to the same very low level in all meal schedule conditions in S1, completely erasing the previously observed anticipatory effect. Conversely, exenatide induced hypoglycaemia in an anticipatory meal schedule dependent pattern. Interestingly, the previously observed positive contrast effect was spared in S2, with a weaker residual effect specifically on the consumption of the more palatable pellet type. To conclude, the food intake reducing effects of exenatide may temporally evolve from strong anorectic to weak anhedonic modulations, where hedonic experience and anticipation during the early anorectic phase is conserved but uncoupled from food intake behaviour.

## Introduction

In general, food intake (what, when, and how much food is consumed) is controlled by the interaction between biological and environmental factors and is modulated by multiple neuronal pathways controlling homeostasis, reward processing and decision making. In addition, to best serve survival, energy homeostasis is directly regulated by overlapping neuronal and hormonal signalling that alters appetite, food craving and foraging behaviour to balance short-term fluctuations in energy intake. High-palatability food can enhance its own intake even beyond homeostatic levels via hedonic sensitization [1], thus leading to regular overeating and hence to obesity [2].

Specific gut hormones, such as glucagon-like peptide 1 (GLP-1) are released in response to ingestion and regulate food intake both in animals and humans by influencing specific appetite-regulating circuits in the brain. GLP-1 plays a role in both the homeostatic and hedonic control of feeding behaviour [3]. GLP-1 is a member of the incretin neuropeptide family, secreted by intestinal enteroendocrine L-cells of the distal ileum and large intestine [4,5] and also in the nucleus of the solitary tract (NTS) in response to nutrient intake [6,7]. Under normal physiological conditions, GLP-1 receptors (GLP-1R) are expressed on the dendritic terminals of vagal afferents innervating the gastrointestinal tract [8], in diencephalic brain regions involved in homeostatic and metabolic control of feeding such as the lateral hypothalamus, and the nucleus arcuatus, and also in other brain regions that specifically regulate reward processing and motivated behaviour such as the ventral tegmental area (VTA) and the nucleus accumbens (NAc) [9]. Among its various actions, GLP-1 stimulates glucose-dependent insulin secretion, inhibits glucagon secretion, delays gastric emptying, enhances feelings of satiety, and consequently reduces food intake [10–12]. In addition, GLP-1 has also been demonstrated to play a major role in maintaining glucose homeostasis by lowering blood glucose levels specifically when the concentration of blood glucose exceeds the normal fasting levels [13].

Exenatide (Ex), a 39-amino-acid GLP-1 analogue peptide, is a potent agonist with high affinity for GLP-1Rs and has a significantly longer biological half-life than the exogenously administered GLP-1, which makes it an efficient treatment option in noninsulin-dependent diabetes mellitus [14–17]. Exenatide reduces postprandial and fasting plasma glucose levels via an insulinotropic effect and via the suppression of plasma glucagon levels in the fasted state [18]. In addition, Ex also penetrates the blood-brain barrier and acts on central GLP-1Rs [19]. In accordance with the behavioural consequences of the ensuing widespread central and peripheral effects of GLP1-R activation, Ex strongly reduces appetite and food intake in rodents, non-human primates, and humans likewise [3,20–22].

Data from animal studies further suggest that the appetite suppressant (anorectic) effects of GLP-1R agonists are also manifested through mesolimbic GLP-1R activation that, in turn, alters the rewarding value of food [23–29]. These effects are especially important with respect to treatment of hedonic overeating in obesity, a more recent indication of GLP1-R agonists [30,31]. The hedonic value of a food is determined by its palatability (flavour, texture, etc.), the environment in which the food interacts with the subject, and ultimately the homeostatic and motivational state of the subject. Motivational states are modulated by specific anticipatory and decision-making cognitive domains, which have a fundamental role in the regulation of the intake of foods that are desired or craved, especially in non-human primates and humans. Despite the success of GLP-1Rs in targeting hedonic overeating in obesity, the way homeostatic, hedonic, cognitive and contextual factors are entwined in the regulation of eating in humans remains unclear [32]. For example, human diets are organized around specific habitual and planned meals scheduled around a day, so our food consumption is also generally regulated by anticipation of a possible later availability of a food item, and also the hedonic value of the available and anticipated food. While relevant research in rodents have successfully encapsulated similar phenomena in clever experimental designs [33], in this domain the differences between rodents and humans are arguably larger than in the case of homeostatic regulation serving basic survival instincts. To increase the translational potential of the available preclinical models, it has become necessary to devise humanized meal-schedule tasks in non-human primates and test the effects GLP1-R modulation on primate scheduled food intake, especially as only a limited number of studies are currently available to support the existing clinical findings and foster further research in the field.

To address this gap, we previously developed a two-session operant food intake paradigm with four meal schedule conditions: two types of pellets with different palatability values were offered as meal in all combinations with two daily feeding sessions (S1 and S2) [34]. In this paradigm, the total daily food consumption of lean and healthy adult animals remained stable over several weeks on a level that clearly avoided overconsumption and covered their homeostatic needs [34]. Within these homeostatic limits, animals consumed more of the more palatable pellet offered in S2 after an S1 with the less palatable meal– a marked successive positive palatability contrast effect. More surprisingly, in the previous experiments, meal schedules were assigned on a weekly basis, and after the current weekly meal schedule had been revealed to the animals on Mondays (diet naïve state), the food consumption pattern of the animals changed in a remarkable way on Tuesdays and remained so for the rest of the week (diet-aware state). Specifically, the animals displayed a strong, palatability-driven negative anticipatory effect in S1: food consumption strongly decreased in anticipation of the more palatable S2 meal, compared to when the less palatable meal was offered in S2.

In the present study, utilizing the above-described novel, highly translatable two-session operant food intake paradigm, rhesus macaques were administered GLP-1R agonist Ex to elucidate how GLP-1R agonism differentially modulates homeostatic feeding regulation, hedonic valuation and palatability-driven anticipation and decision making across the two daily feeding sessions.

## Materials and Methods

### Animals and housing

Five 4-year-old, pair-housed, pre-trained, healthy male rhesus macaques (*Macaca mulatta*) were used in the experiments. Their body weight was 4.4±0.3 kg and 5.5±0.5 kg (mean±SEM) at the beginning and at the end of the study, respectively. The animals were kept in home cages that were uniformly sized 200 × 100 × 200 cm (length × width × depth) and were equipped with wooden rest areas. Besides natural light from windows, illumination from full-spectrum artificial light was provided for 12 hours starting at 7:00 AM each day. Temperature (24±1°C) and relative humidity (55±10%) of the animal house were kept constant, with continuous airflow.

On experimental days (weekdays, Mondays to Fridays), daily food allowance was only offered during the food intake sessions (see section ‘Two-session operant food intake paradigm’ below). On weekend days (Saturdays and Sundays), animals received usual dry lab chow (Altromin Spezialfutter GmbH & Co, Lage, DE; nutritionally complete, 3.315 kcal/g) with a variety of vegetable and fruit supplements. The calorie intake from the pellets consumed during the experimental days and calorie intake on weekend days were similar, and fully complied with the recommended daily energy requirements of the animals (see section ‘Energy requirements’ in **Supplementary Methods**). Water was *ad libitum* available on every day of the week.

### Ethical compliance

All procedures were conducted in the Grastyán Translational Research Centre of the University of Pécs. The study was approved by the Animal Welfare Committee of the University of Pécs, and the Hungarian National Scientific Ethical Committee on Animal Experimentation. Ethical permission was issued by the Department of Animal Health and Food Control of the County Government Offices of the Ministry of Agriculture (permission number: BA02/2000-13/2015). Measures were taken to minimize pain and discomfort of the animals in accordance with the Directive 40/2013 (II.14): “On animal experiments” issued by the Government of Hungary, and the Directive 2010/63/EU “On the protection of animals used for scientific purposes” issued by the European Parliament and the European Council.

### Operant chambers

Training and experimental sessions were conducted in individual, isolated, ventilated and illuminated primate operant conditioning chambers which were located in a separate testing room next to the animal house. Each operant chamber was equipped with a response panel (Modular Intelligence Test System for Primates, Med Associates, St. Albans, Vermont, US) and a built-in standard digital camera for real-time video surveillance. Operant response panels were controlled with the MED PC software for MS Windows (Med Associates, St. Albans, Vermont, US).

### Two-session operant food intake paradigm

The experiments were performed using a two-session operant food intake behavioural paradigm that was developed in our laboratory [34]. All the data analysed in this study were collected on the experimental weeks when all animals were involved in the two-session operant food intake paradigm on each weekday. Sessions consisted of repeated individual trials where the animals received a single one-gram food pellet upon three consecutive lever presses (fixed ratio 3 reinforcement schedule, FR3) per trial. At the beginning of each trial, a cue light was switched on, and was lit until the three lever presses were completed (without any time limit), which was followed by the delivery of the reward pellet and a 30-sec inter-trial interval (ITI, see **Figure 1C**). Lever presses during the ITI were neither rewarded with food pellets nor punished with extra time-out. Two experimental sessions were run per day: Session 1 (S1) started at approx. 9:30 AM and lasted 2 h (or until max. 200 pellets were earned), and Session 2 (S2) started at approx. 12:30 PM and lasted 1 h (or until max. 100 pellets were earned). There was a one-hour inter-session interval (ISI) between S1 and S2 when the subjects were returned to their home cages (see **Figure 1B**).

**Figure 1.**
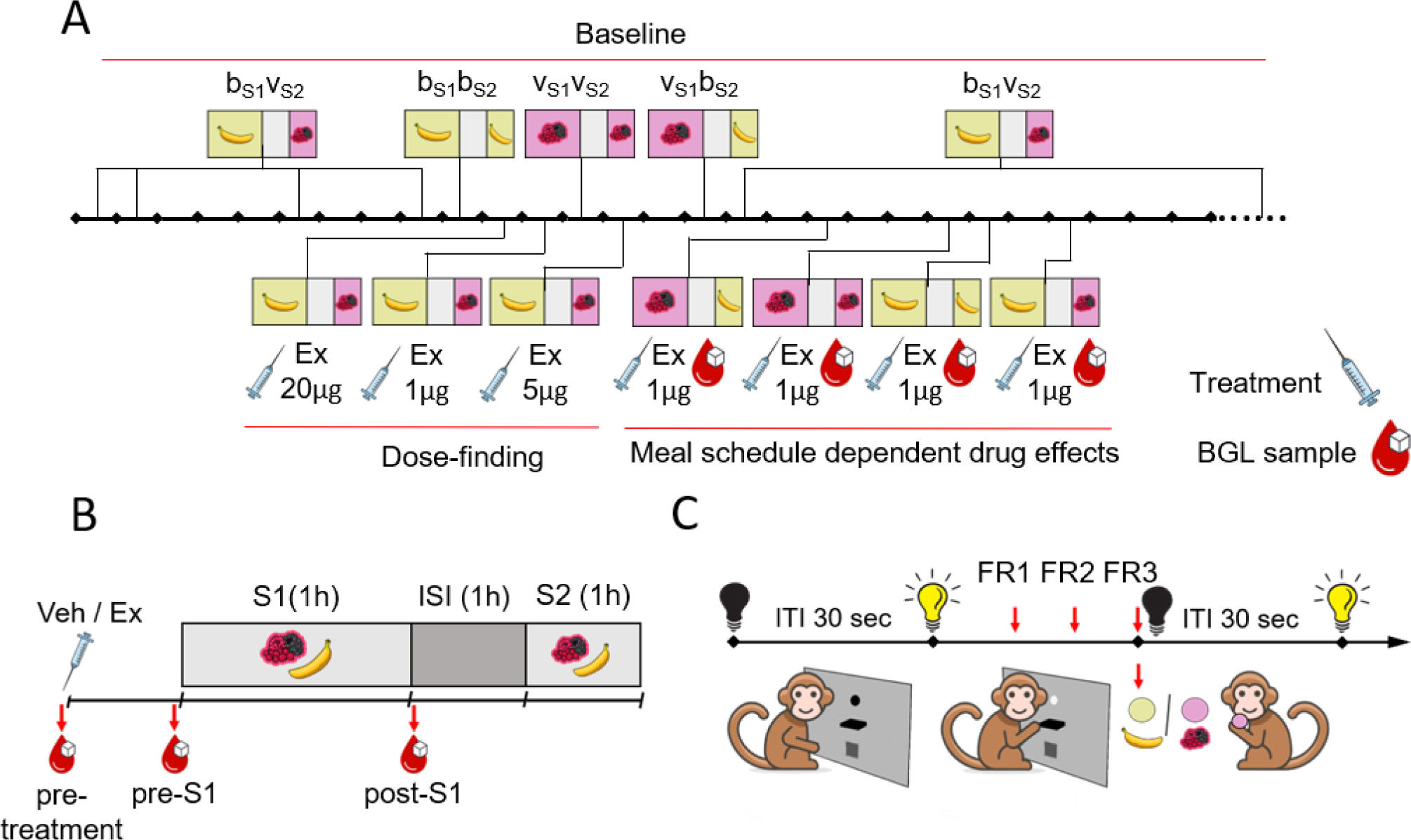
Timeline and schematic experimental design. **(A) Timeline of the experimental weeks.** On the timeline, pictograms and acronyms indicate the weekly meal schedule for all experimental weeks. Experimental weeks without pharmacological treatments are called Baseline weeks as indicated above the timeline. Below the timeline, the treatment weeks of the *Dose-finding* and *Meal schedule experiments* are shown, labelled with treatment types (syringe pictogram), administered doses, and blood sampling (blood drop pictogram). Treatment weeks involved both Vehicle and Exenatide treatment days (see panel B and main text). **(B) Daily schedule of the food intake task.** On each weekday the animals performed the food intake task for 2 experimental sessions (S1 and S2) separated by a one-hour intersession interval (ISI). Within a session, only one type of food pellet was offered: either less palatable (banana flavoured) pellets or more palatable (‘very berry’ flavoured) pellets, as also indicated by pictograms. Timing of treatment administration and blood sampling are indicated by pictograms. **(C) Trial structure of the food intake task.** At the beginning of each trial, a cue light switched on, indicating that food items were available upon three consecutive lever presses (red arrows, fixed ratio 3 reinforcement schedule, FR3). After pellet delivery, the cue light was switched off for a 30-sec intertrial interval (ITI).

The session durations and the number of available pellets were previously determined so as not to limit the animals’ food consumption: There were only 3 out of 175 occasions for S1 and 9 out of 175 occasions for S2 when the sessions were 1-5 minutes shorter because of reaching the maximal pellet consumption.

During a session, food pellets of only one type were available: either the standard, low-palatability banana flavoured pellets (denoted ‘b’, Dustless Precision Pellets^®^, Bio-Serv, Inc., Frenchtown, New Jersey, US; nutritionally complete, 3.35 kcal/g) and the more palatable ‘very berry’ (forest fruit) flavoured pellets (denoted ‘v’, Supreme Mini-Treats, Bio-Serv, Inc., Frenchtown, New Jersey, US; nutritionally complete, 3.46 kcal/g). Food pellet types were selected based on food preference test, as described in our previous study [34]. All possible combinations of the two types of pellets assigned to the two daily sessions were tested, resulting in 4 different meal schedule conditions: S1 banana / S2 banana (b_S1_b_S2_), S1 banana / S2 very berry (b_S1_v_S2_), S1 very berry / S2 banana (v_S1_b_S2_), S1 very berry / S2 very berry (v_S1_v_S2_). Each different meal schedule was consistently offered for 5 consecutive experimental days, from Mondays to Fridays.

### Procedures and drug administration

On Baseline weeks we recorded the animals’ feeding behaviour in the two-session operant paradigm without pharmacological treatments. Two experiments were conducted, a *Dose-finding experiment* involving 3 treatment weeks with the b_S1_v_S2_ meal schedule, and the *Meal schedule experiment* involving 4 treatment weeks, one for each of the 4 meal schedule conditions. On treatment weeks, Vehicle treatment was administered on Tuesdays, and drug treatment was administered on Wednesdays. The timeline of all the experimental weeks – treatment and Baseline weeks is shown on **Figure 1A**.

We investigated the effects of peripheral administration the glucagon-like peptide-1 agonist exenatide (Ex, Byetta 10 µg injection pen; synthetic exendin-4; AstraZeneca AB., Stockholm Sweden; batch number: 15H029). In our experiments, animals received acute, subcutaneous (under the skin of the thigh) injection of Ex on Wednesdays, 1 hour prior to S1. Exenatide was extracted from the injection pen and was diluted with a vehicle solution (pH=4.5, acetate buffer with mannitol) containing the main components of the original vehicle from the product to achieve a larger dosing volume (0.1 mL/kg) set based on individual body weight. Exenatide or vehicle solutions were administered via single subcutaneous injections. The daily dosing formulations were prepared freshly (max. 2 hours) prior to administration. The vehicle solution was also used for Vehicle (sham) treatments which were administered on Tuesdays with the same timing as the genuine treatments. At least 6 days of recovery time was introduced between pharmacological treatments to eliminate possible residual effects from the previous treatment and to enable monitoring food intake on subsequent days (Thursdays and Fridays).

In the *Dose-finding experiment*, we determined the effective and side effect-free dosing by administering Ex at doses of 1, 5, or 20 µg/kg in meal schedule b_S1_v_S2_ (see **Figure 1A**). All subjects were observed and were assessed for side effects such as changes in alertness and general activity and the presence of excessive salivation, gagging, and regurgitation following each injection until the end of experimental day (4 pm). Following the *Dose-finding experiment*, we conducted a *Meal schedule experiment*, where we investigated the acute effects of 1 µg/kg Ex in all four meal schedules. The dosing formulation protocol, the route of administration, dose volumes, and the timing of dosing were the same in both of experiments.

### Blood glucose level sampling

For the measurement of blood glucose levels (BGL) in the *Meal schedule experiment*, blood samples were taken from the monkeys seated in a standard primate chair, which was otherwise used for the transport of animals and for experimental sessions from their home cages. Monkeys had been previously accustomed to the procedure of taking blood samples and trained to voluntarily offer their hind limb for sampling to minimize stress. After puncturing the skin with a needle tip on the large toe of the hind limb, a drop of blood was applied to the test strip. Blood glucose levels were measured with EasyTouch blood glucose monitor (Wellmed EasyTouch® GCU, ET-301 Multi-Function Monitoring System, Bioptik Technology Inc., Jhunan Township, TW) in all four meal schedule conditions according to the following timetable: immediately before administration of drug or vehicle (pre-treatment); just before S1 (pre-S1) and after S1 (post-S1).

### Statistical Analysis

Data loading and preparation was partly done in MATLAB R2021a (The Mathworks, Natick, MA, USA); analyses and graphics were made in R 4.3.2. Where applicable, statistical tests are two-sided.

#### Food consumption analyses

We investigated the total food consumption (g) for each session. The number of pellets delivered was registered by the operant testing apparatus. Leftover pellets were collected from the operant chambers at the end of the sessions and their number was registered manually. Food consumption for each session was calculated by subtracting the number of leftover pellets from the number of pellets delivered.

For both experiments, two different control conditions were used: Baseline and Vehicle control. Specifically, for the *Dose-finding experiment*, the Vehicle control was data averaged from the 3 Vehicle sessions from the days (Tuesdays) preceding the drug administration days. Baseline control for the *Dose-finding experiment* was the average from 6 sessions from the same days of the Baseline weeks as the drug administration days (Wednesdays). We verified using linear and quadratic contrasts in repeated measures ANOVAs that there were no significant trends across Baseline and Vehicle control days across the weeks (see ‘Dose finding experiment’ in **Supplementary Results**). Vehicle controls for the *Meal schedule experiment* were not averaged, one Vehicle day from the same week was used for each (weekly) meal schedule condition. Baseline control for the analysis of the *Meal schedule experiment* was calculated using days from Tuesday to Friday averaged, from Baseline weeks with the corresponding meal schedule conditions. The Baseline control weeks of the *Meal schedule experiment* were analysed in detail in our previous paper where we showed that the animals displayed a clear anticipation effect from Tuesdays to Fridays [34]. Since our present aim was to investigate the pharmacological modulation of this effect, we deliberately included Tuesdays to Fridays (diet-aware days) in the experimental design and analysis (see also the Introduction).

Data analyses for S1 and S2 were conducted separately as, due to their differing duration, food consumption levels and corresponding variances were different between the two sessions. Thus, for both experiments, four repeated measures ANOVAs were conducted on food consumption data, separate ones for the two control conditions used and for S1 and S2 (S1 with Vehicle control, S1 with Baseline control, S2 with Vehicle control, S2 with Baseline control).

The ANOVAs for the *Dose-finding experiment* involved only the Treatment factor (4 levels: Control – Vehicle or Baseline, Ex 1 µg/kg, Ex 5 µg/kg, Ex 20 µg/kg). The ANOVAs for the analysis of the *Meal schedule experiment* involved the following factors: S1 meal type (b_S1_ or v_S1_), S2 meal type (b_S2_ or v_S2_) and Treatment (2 levels: either Vehicle or Baseline control, Ex 1 µg/kg).

#### Daily energy intake analysis

The daily energy intake was calculated based on the food consumption values described above and the ME content of the pellets as supplied by the manufacturer. Repeated measures ANOVAs identical to those performed on S1 and S2 were run on daily energy intake data, except that the models were parametrized so that each animal’s recommended daily energy intake (see ‘Energy requirements’ in Supplementary Methods) was the intercept. This way we could also quantify and statistically test the difference of each Treatment condition (including the Baseline and Vehicle controls) from the daily nutritional recommendation. Since the main focus of this analysis was relating the recommended daily intake, further results were considered redundant with the detailed S1 and S2 analyses and are not reported here.

#### Blood glucose level analysis

We analysed the BGL data from samples taken during the *Meal schedule experiment*. Please refer to the section ‘Blood glucose level analysis’ in the **Supplementary Methods** for details.

## Results

We first conducted a *Dose-finding experiment* using meal schedule b_S1_v_S2_ to probe the effects of Ex on food intake in the two-session operant paradigm. Based on the results, we selected the optimal dose of Ex which reduced food intake without observable side effects, and went on to investigate the hypothesized meal schedule dependent effects of Ex (see **Figure 1**).

### Dose-finding experiment

In the *Dose-finding* experiment the total daily energy intake significantly and dose-dependently decreased under Ex treatment (F_2.0,8.1_=60.21, p=1.3×10^−5^ using the placebo-treatment Vehicle control; F_1.8,7.2_=38.24, p=0.0002 using the no-treatment Baseline control; see **Figure 2B**). The Vehicle and Baseline control energy intake did not differ significantly from nutritional recommendations based on age and body weight, but the daily energy intake was significantly lower than Vehicle, Baseline, and also the recommended nutritional value for the 5 and 20 μg/kg doses of Ex, while it only marginally decreased for the 1 μg/kg dose (see **Supplementary Table 1** for details).

**Figure 2.**
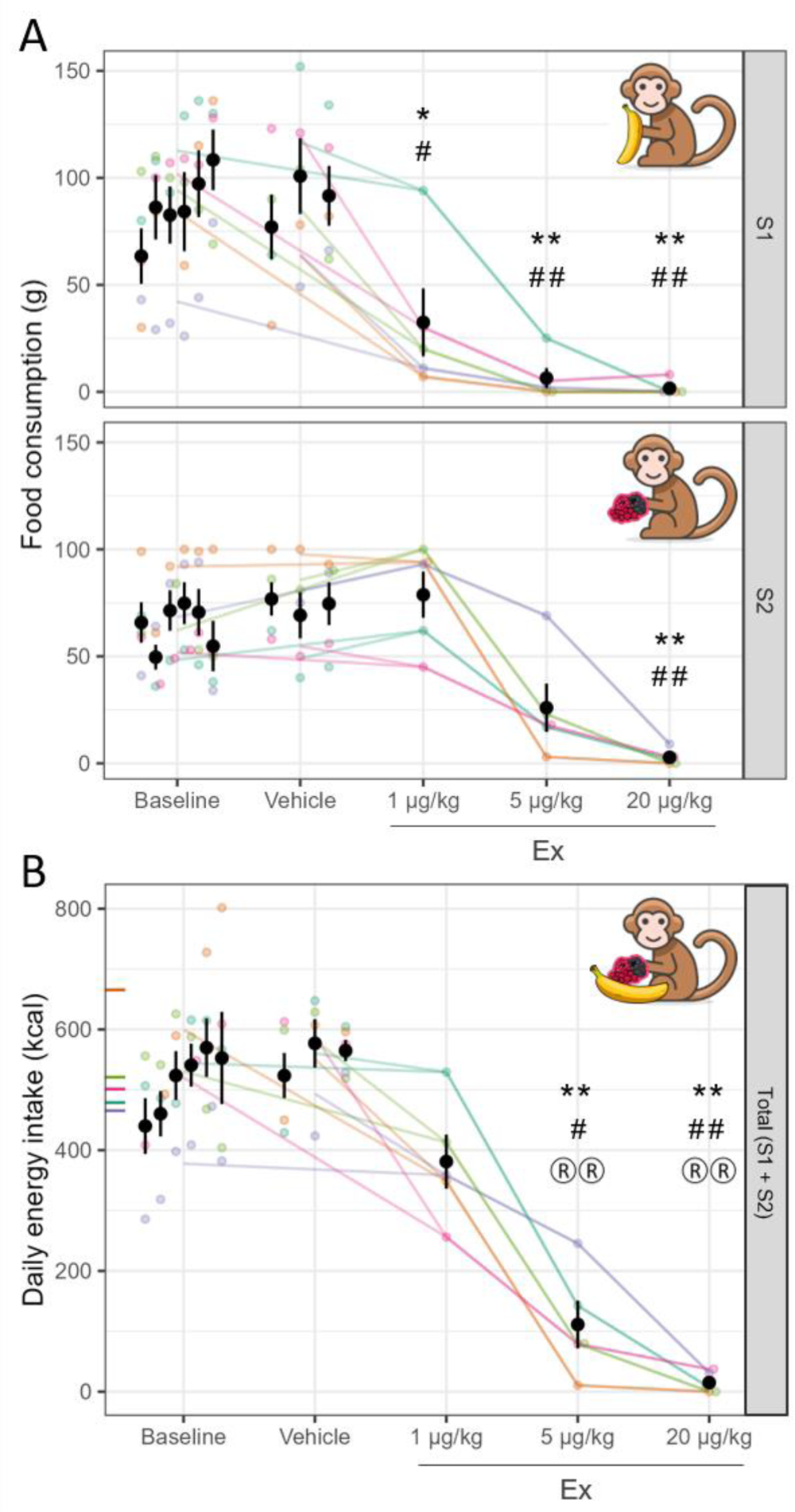
Dose-finding experiment in the food intake paradigm. (A) Dose-dependent effects of Ex on food consumption and (B) on total daily energy intake. Effects of 1, 5 or 20 µg/kg Ex (x axis) on food consumption (y axis on A) in S1 (upper panel) and S2 (lower panel) and daily energy intake (y axis on B) of the two-session food intake paradigm compared to Vehicle control days (Vehicle, *: p<0.05,**: p<0.01) and no-treatment Baseline days (Baseline, #: p<0.05, ##: p<0.01) p-values are Dunnett corrected for 3 comparisons per session. Colors indicate individual animals, points correspond to data from one session on panel A and one day on panel B. Black points and whiskers are condition averages and s.e.m across animals, with the 6 Baseline and 3 Vehicle days shown separately. On the y axis of panel B, inward pointing ticks show the daily energy intake recommendation for each animal (color-coded) based on their current weight. On panel B, ® markers denote significant differences from nutritional recommendation values (®: p<0.05, ®®: p<0.01).

In S1, Ex significantly decreased food consumption in all three doses, regardless of which control condition was used as reference (see **Figure 2A**, top **Supplementary Table 2**, top), with close to zero consumption in the two higher (5 and 20 µg/kg) dose levels. In S2 overall, Ex still caused strong, significant decrements in food consumption (see **Figure 2A**, bottom; **Supplementary Table 2**, bottom), however, the 1 µg/kg dose of Ex was not effective, and close-to-zero floor consumption levels were only reached for the highest (20 µg/kg) dose of Ex.

In addition to effectiveness, in the case of the 20 µg/kg dose of Ex, various side effects were observed. All of the animals showed transient hypoactivity, coordination problems and gastrointestinal symptoms. After the administration of the 5 µg/kg dose of Ex, side effects were observed only in one animal, while in the case of 1 µg/kg dose, no side effects were observed, all animals showed normal behaviour and general activity.

### Meal schedule dependent experiment

We went on to investigate the effects of the 1 µg/kg dose of Ex on food consumption in the complete version of our two-session operant food intake paradigm in all four possible meal schedule conditions (see **Figure 3**).

**Figure 3.**
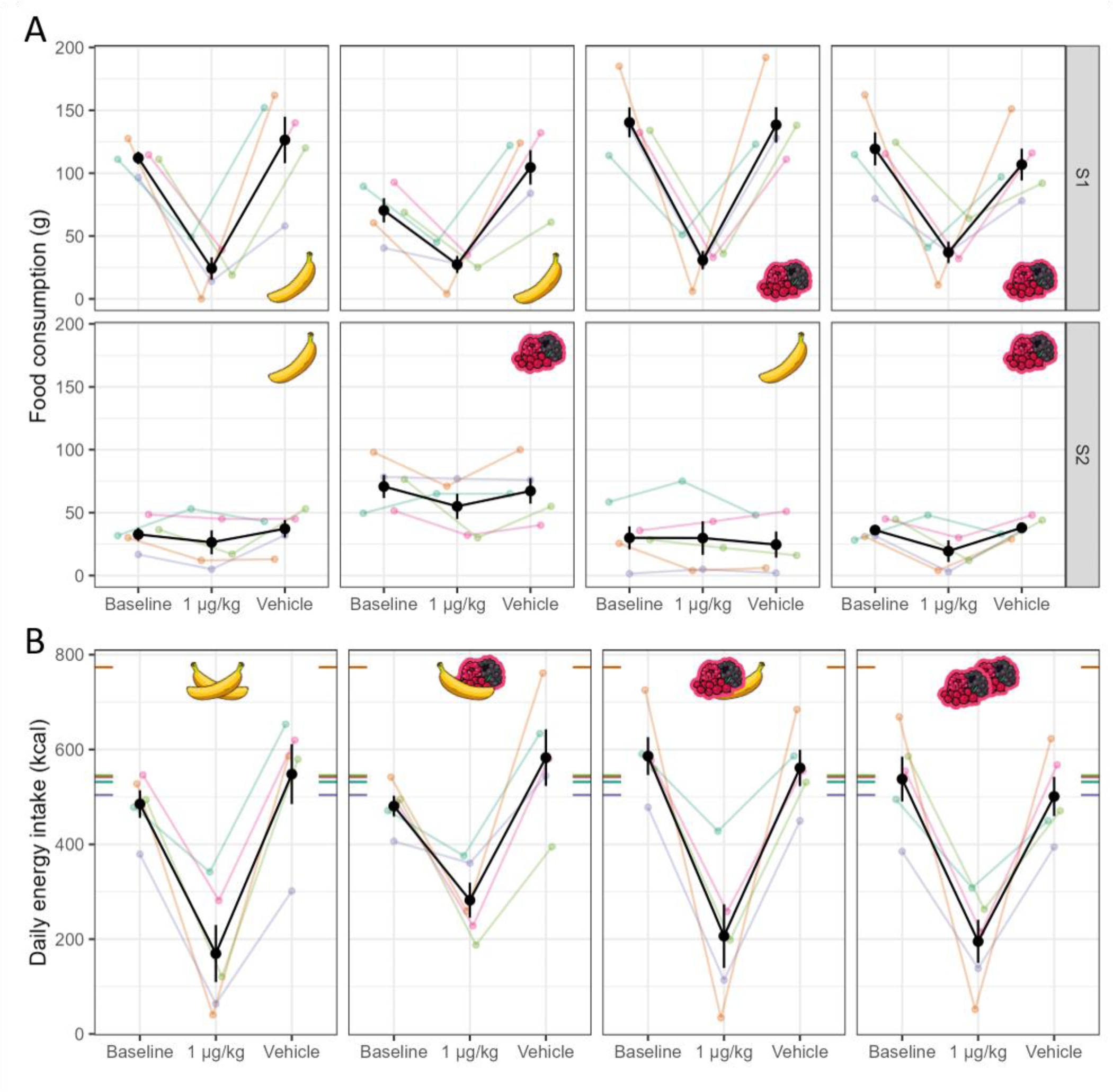
Meal schedule experiment in the food intake paradigm. The effects of 1 µg/kg Ex across meal schedule (panels from left to right) and treatment conditions (x axis) on food consumption (y axis, panel A) across sessions (lower and upper part of panel A), and on daily total energy intake (y axis, panel B). Pictograms represent meal schedule conditions Colours indicate individual animals; coloured points correspond to data from one session on panel A and one day on panel B. Black points and whiskers are condition averages and s.e.m across animals. Coloured tick marks on the edges of sub-panels on panel B show recommended energy intake of each animal calculated based on weight measured during the experiment.

Total daily energy intake significantly decreased under 1 µg/kg dose of Ex treatment relative to nutritional recommendations (see **Figure 3B**, d=−366 ± 84 kcal, t_4_=−4.35, p=0.012), Baseline (d=−309 ± 62 kcal, t_4_=−4.99, p=0.0075) and Vehicle controls (d=−335 ± 62 kcal, t_4_=−5.45, p=0.0055). In contrast, energy intake in the Baseline and Vehicle conditions matched the nutritional recommendations (relative to Baseline: d=−57 ± 30 kcal, t_4_=−1.91, p=0.13; relative to Vehicle: d=−31 ± 32 kcal, t_4_=−0.97, p=0.38).

In our previous study [34] validating and describing this paradigm, we found a palatability-driven anticipation effect: S1 food consumption in the two-session paradigm strongly decreased in the b_S1_v_S2_ condition in anticipation of the more palatable S2 meal type, compared to when the less palatable meal was offered in S2 (the b_S1_b_S2_ condition). Here, this effect can be seen in the Baseline condition (Baseline, S2 meal type: F_1,4_=13.80, p=0.021, see also Figure 3). (Note that the Baseline control data has been adopted from our previous study [34] to facilitate comparison - see also Methods). Analyses with Vehicle control days yielded comparable results, most of these are shown in the respective sections of the **Supplementary Results**.) S1 food consumption was also boosted by the palatability of the currently consumed food (Baseline, S1 meal type: F_1,4_=11.35, p=0.028), but the interaction of these two effects was not significant (Baseline, S1 meal type × S2 meal type: F_1,4_=5.95, p=0.071).

Under Ex treatment, food intake in S1 strongly decreased (main effect of Treatment: F_1,4_=41.96, p=0.003) to the same low level (from 110.5 ± 7.3 g in Baseline to 29.9 ± 6.9 g in Ex) regardless of meal schedule condition (under Ex treatment, S1 meal type: F_1,4_=2.01, p=0.23; S2 meal type: F_1,4_=1.14, p=0.35; S1 meal type × S2 meal type: F_1,4_=0.35, p=0.58), as also manifested in the Treatment × S1 meal type (F_1,4_=7.57, p=0.051) and Treatment × S2 meal type (F_1,4_=9.72, p=0.036) interactions. Thus, the palatability-driven anticipatory effect in S1 was completely erased by the 1 µg/kg dose of Ex.

The characteristic pattern in S2 was the successive positive palatability contrast effect – a dramatic increase in food intake in the b_S1_v_S2_ meal schedule condition compared to all the remaining three meal schedule conditions. Importantly, this pattern was fully preserved under Ex treatment: it generalised across the control conditions and the treatment condition as demonstrated by the significant S1 meal type × S2 meal type interaction (using Baseline days as control: F_1,4_=9.53, p=0.037) not moderated by Treatment (Treatment × S1 meal type × S2 meal type using Baseline days: F_1,4_=0.60, p=0.48).

Besides leaving the successive positive contrast effect in S2 unaffected, Ex had a slightly stronger effect on the consumption of the more palatable food (mean ± SE average treatment effect: −16.2 ± 9.5 g against Baseline and −15.4 ± 6.5 g against Vehicle) compared to the less palatable food (−3.2 ± 6.7 g against Baseline and −2.8 ± 5.9 g against Vehicle) as supported by a Treatment × S2 meal type interaction (using Baseline days as control: F_1,4_=8.46, p=0.044, using Vehicle days as control: F_1,4_=12.37, p=0.025).

Blood glucose levels (BGL) were measured during the four meal schedules both on the Vehicle and the Ex treatment days (see **Figure 4**). BGL in general substantially dropped from pre-treatment to pre-S1 (F_1,4_=8.79, p=0.041, change-score analysis – further analyses are baseline-adjusted models or raw values. (For more detailed statistics, see **Supplementary Results**). During Vehicle treatment in S1 with very berry flavoured pellets offered, BGL increased (from pre-S1 to post-S1) by 0.76 ± 0.13 mmol/L, a significantly (Time × S1 meal type: t_50.0_=4.30, p=8×10^−5^) larger increase compared to sessions when banana flavoured pellets were offered, where it stagnated (+0.06 ± 0.13 mmol/L). This effect was erased by Ex treatment (post-S1 minus pre-S1, very berry consumed, Ex treatment: +0.11 ± 0.18 mmol/L), which is not surprising given the similar low consumption rates across conditions under the effect of the drug.

**Figure 4.**
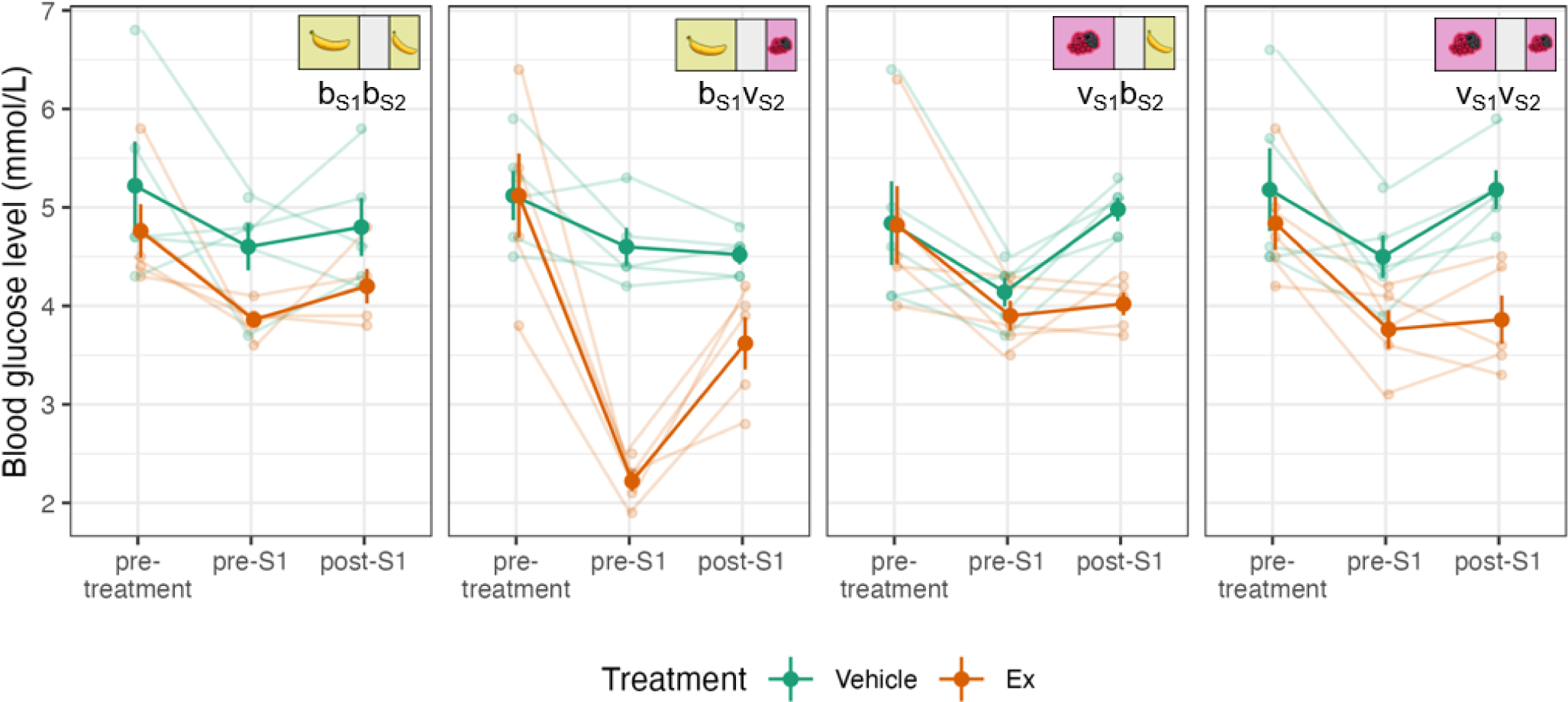
Blood glucose levels in the Meal schedule experiment. Blood glucose levels (y axis) from the three sampling times (pre-treatment, pre-S1, post-S1, on x axis) in the four meal schedule conditions (panels left to right) under Vehicle and Ex treatment (green and orange, respectively).

Exenatide strongly decreased pre-S1 BGL specifically in the b_S1_v_S2_ meal schedule condition (d=−2.38 ± 0.25 mmol/L, t_6.8_=−9.7, p=3×10^−5^, uncorrected contrast), and only to smaller extent in the other conditions (d ≥ −0.73, t ≥ −3.5, p ≥ 0.006; pre-S1, Treatment: F_1,4.2_=41.50, p=0.0025; Treatment × S1 meal type × S2 meal type: t_50.4_=3.50, p=0.001). This effect substantially weakened in the post-S1 blood sample (Treatment × Time × S1 meal type × S2 meal type interaction in the main model: t_50.0_=−2.61, p=0.012), though BGL was still significantly lower under Ex compared to Vehicle control treatment (post-S1, Treatment: F_1,4.1_=24.23, p=0.0073).

## Discussion

In the present study, we investigated the potential effects of Ex on palatability-based food preference and anticipation in rhesus macaques, besides confirming its known effects on homeostatic food intake regulation. In the *Meal schedule experiment*, we found that 1 µg/kg dose Ex treatment strongly decreased food consumption in S1 (1 to 3 hours after treatment) and erased the previously observed palatability-driven negative anticipatory effect. Interestingly, no marked food intake reduction effect of Ex was observed in S2 (4 to 5 hours after treatment), and, surprisingly, the previously observed successive positive contrast in food intake in S2 also persisted under Ex treatment. However, Ex treatment was shown to mildly reduce intake of the more palatable, but not the less palatable food in S2. Additionally, Ex decreased both fasting and postprandial blood glucose levels (BGL) measured before and after S1, and the Ex effect on fasting BGL depended on the meal schedule.

Extending our previous results, here we confirmed that in control conditions, homeostatic regulation of food consumption was captured by the two-session feeding paradigm as the daily energy intake in the operant task matched nutritional recommendations to maintain growth and energy homeostasis. Ex, in line with its established effect on food intake (REF), dose-dependently decreased food consumption in our paradigm so that the energy intake fell below both control and recommended nutritional values.

Though no studies have analysed the anorectic effects of Ex in non-human primates so far in a paradigm with repeated, scheduled feeding sessions, the previous results of Scott and Moran on rhesus macaques are still broadly compatible with our present findings [22]. In their study [22], animals were tested in a single six-hour operant food intake session starting 15 minutes after intramuscular administration of Ex. Despite important procedural differences from our present study, a closer look at their findings also reveals a strong initial anorectic effect of Ex in the first 1to 3 hours, and the food consumption reductions in the 4 to 6-hour time period were of comparable size to our present findings in S2. We extended those earlier results by investigating the effect of Ex under different meal schedules, showing that the late effect of Ex is not only smaller but depends on the palatability of the offered food, implying that it is also more context-dependent compared to the strong early effect of Ex.

In the Baseline condition, when the more palatable food was available in S2, the animals reduced their consumption in S1 anticipating the more valued meal, so that the consumption decrement in S1 matched the excess amount eaten in S2 – we referred to this phenomenon as the palatability-driven negative anticipatory effect [34]. In the *Meal schedule experiment*, after the acute administration of the 1 µg/kg dose of Ex food consumption decreased to the same very low level in all four meal schedule conditions, that is, the palatability effect and the palatability-driven negative anticipatory effect were both erased. These results suggest that the early effect of 1 µg/kg Ex decreases the homeostatic drive, i.e., hunger, to a level so low that is known to weaken or eradicate hedonic contrast effects [33,35,36].

In S2, we did not observe a strong, meal schedule independent reduction in feeding that would have been expected given a sustained pharmacological effect, but neither did we find compensatory consumption after the reduced S1 feeding rates that would have been expected due to declining drug effects. Food consumption in each condition was approximately in the same range as that of the respective control (no-treatment Baseline or placebo-treatment Vehicle) conditions with the same meal schedule. Importantly, the characteristic successive positive palatability contrast effect, that is, the excess S2 consumption on the b_S1_v_S2_ meal schedule condition relative to all other conditions, was also conserved. This positive contrast effect requires awareness of the hedonic values of both the presently experienced and the previously consumed food types. Based on this, we hypothesize that even after radically lower food consumption in S1 following the administration of Ex, before entering S2, the hedonic state (or sensory-specific satiation [37]) attained during S1 and carried over to S2 was similar to that of the control conditions.

The positive contrast in S2 was spared by the late effect of Ex. However, Ex specifically decreased the consumption of the more palatable, but not the less palatable food in S2. This is in line with previous studies in rodents showing that systemically administered Ex diminished the previously established palatability-based place preference and general motivation measured by the progressive ratio operant test. Exenatide also decreased preference of sucrose solution against water and more palatable against less palatable food, which are interpreted as anhedonic effects. [24,29,38–41]. Exenatide also had strong anhedonic effects in the same paradigms when injected directly into reward-related brain areas such as the ventral tegmental area (VTA), the nucleus accumbens (NAc), and even in the nucleus tractus solitarii (NTS), a structure previously thought to regulate homeostatic feeding only [25–29]. Results from a blood-oxygen dependent functional MRI study [42] also suggest that mesolimbic reward-related areas may also mediate the anorectic effects of Ex in humans, with a hypothesized mechanism of decreasing responsiveness to contextual food cues (incentive salience, ‘wanting’) and increasing activity that correlates with consummatory reward value (‘liking’) [43]. Thus, GLP-1R activation in reward-related mesolimbic and brainstem nuclei reduces the motivational incentive of the available most rewarding stimulus/food, mediating the hedonic aspects of feeding regulation such as the rewarding value of the palatable food.

The BGL values measured in the pre-treatment samples and on Vehicle control days were consistent with the BGL under laboratory conditions during everyday procedures reported in the literature [44]. In line with clinical and experimental data [45,46], Ex strongly decreased BGL relative to the Vehicle condition both before and after S1. In the case of Vehicle treatment, higher values of postprandial BGL were obtained after S1 when the animals consumed a slightly higher amount of food because the more palatable meal was available in S1. This discrepancy disappeared due to Ex treatment which can be explained by the similar low food consumption in all four meal schedule conditions. Under Vehicle treatment, fasting BGL values consistently decreased during the hour preceding the start of S1. This is in accordance with previous human studies showing that food expectancy and delaying meals lead to decreasing blood glucose levels [45,46]. During food expectation – that is, in the cephalic phase – even thinking about the food can initiate several physiological processes like increased insulin secretion, increased ghrelin levels, increased digestive juice secretion, and also decreasing blood glucose levels [45,47,48]. These processes constitute the early activation of the processes of postprandial metabolism, to prepare the organism for the digestion and absorption of the forthcoming food and nutrients [47]. Exenatide treatment caused a further drop in the fasting BGL in all four meal schedule conditions. Importantly, it was the b_S1_v_S2_ meal schedule condition where the BGL decreasing effect of Ex was exceptionally strong, even pushing BGL into the hypoglycaemic range which was not reached under any other conditions. A possible explanation for this strong hypoglycaemic effect is that the palatability-related anticipation effect that we have shown in our previous study [34] is the strongest in the b_S1_v_S2_ meal schedule condition, and the cephalic phase effects of this anticipatory process were even amplified by Ex. This is a striking observation, especially if we consider the fact that the negative anticipatory effect was completely erased from the S1 food consumption behaviour by Ex.

Several factors have limited the investigation of motivation and food reward sensitivity in animals in previous food intake studies. Frequently, animals were on an ad libitum feeding regime outside the scope of the experiments, where postprandial processes were left uncontrolled. Our novel two-session operant food intake paradigm addresses these limitations using a well-controlled feeding regime that permits the investigation of reward-related food intake behaviour without the confounding influence of food available outside the context of the experiment. Our paradigm also did not require food deprivation: in contrast to several previous laboratory-based experiments (e.g., [49,50]), the animals were allowed to eat in an unrestrained and practically ad libitum manner (see **Methods**). Thus, the behavioural pattern of food consumption in this paradigm can be considered endogenously controlled, so it is better suited to investigate close-to-natural feeding regulation of the animals under laboratory circumstances.

It is important to note that we used lean, healthy young adult animals in our studies. Since the regulation of food intake through these complex processes of dynamic interactions between the homeostatic and hedonic control systems and the higher-order regulatory adjustments functioned normally in the young animals used in our studies, which limits the direct applicability of these results to maladaptive eating behaviour or disorders in the regulation of feeding. However, we believe that this paradigm could be well used in the future to study multiple aspects of feeding regulation in obesity models with dysregulated feeding behaviour (i.e., in animals that overeat during ad libitum feeding).

To sum up, we reported here that, in our novel two-session operant food intake paradigm, GLP-1 agonism differentially modulated homeostatic, hedonic and anticipatory aspects of palatability-driven feeding behaviour. Exenatide had a strong early (1 to 3 hours post-administration, S1) homeostatic anorectic effect, and the previously observed palatability-driven negative anticipatory effect in S1 disappeared completely under Ex treatment. Food consumption levels recovered by S2 (4 to 5 hours post-administration), and in this late phase a weak, palatability-dependent food intake reduction was observed – suggesting an anhedonic effect. The positive palatability contrast effects in S2 were largely conserved, suggesting that even though it did not influence food consumption behaviour, hedonic experience in S1 was conserved under the early Ex effect. Intriguingly, by amplifying preprandial blood glucose level decreases, Ex induced hypoglycaemia specifically in the focal condition of the palatability-driven negative anticipatory effect that was eradicated by it on the level of food consumption behaviour.

To conclude, hedonic behavioural patterns following S1 and anticipatory blood sugar level effects preceding S1 together suggest that during S1 hedonic and anticipatory regulatory mechanisms were present under the early Ex effect, but were uncoupled from ongoing feeding behaviour. Understanding the underlying biological pathways and their complex interactions in disease-models will open new avenues in designing more targeted therapies and predicting potential therapeutic outcomes in humans.

## Acknowledgements

We would like to thank Zoltán Gödri for valuable technical contribution and for their assistance in animal care. We would like to thank Dr. Attila Trunk for technical help with experiment setup and initial data processing.

## Author contributions

Conceptualization, BK*, JI*, PK and IH; methodology, BK, JI, PK and IH; Investigation, JI; software, BK; formal analysis, BK; data curation, BK, JI; writing – original draft, BK, JI and IH; writing – review & editing, BK, JI, PK, BL and IH; visualisation, BK, JI; resources, BL and IH; project administration, BL and IH; funding acquisition, BL and IH; supervision, IH.

*These authors contributed equally to this work.

