## Supplementary Information for "GLP-1 receptor agonist exenatide uncouples food intake from hedonic and anticipatory regulation in non-human primates: Insights from an operant meal schedule paradigm"

for

### Supplementary Methods

**Energy Requirements**

Juvenile nonhuman primates require more energy per unit of body weight for growth than do adults of their species. For juvenile rhesus macaques, the daily metabolizable energy (ME) intake requirement was determined as 107 $kcal \cdot BWkg^{-1} \cdot d^{-1}$ [51]. For the subjects participating in our study, the mean daily ME intake requirement during Baseline weeks was 515.3±40.7 $kcal \cdot d^{-1}$ (at the commencement of the study: 468.2±33.8 $kcal \cdot d^{-1}$; at the end of the study: 562.4±48.5 $kcal \cdot d^{-1}$). The daily ME intake, calculated based on the measured food consumption during Baseline weeks and the ME of the pellets supplied by the manufacturer, corresponded to the recommended daily ME intake in all four meal schedules ($b_{S1}b_{S2}$: 485.1±29.0 $kcal \cdot d^{-1}$; $b_{S1}v_{S2}$: 480.6±21.9 $kcal \cdot d^{-1}$; $v_{S1}b_{S2}$: 586.1±39.9 $kcal \cdot d^{-1}$; $v_{S1}v_{S2}$: 560.7±43.8 $kcal \cdot d^{-1}$).

The effect of Exenatide in terms of daily ME intake was also analysed as described in the Statistical analysis section in the Methods section of the main text. Recommended ME intake was calculated based on weights measured on two days from between the first and last week of the experiment.

#### Technical errors during testing

During the entire experimental period, we registered 4 technical errors (on 3 Baseline sessions and 1 Vehicle session, see ‘Procedures and drug administration’ in the main text), caused by the jamming of the pellet dispenser, in these cases, at the end of the sessions involved, we subsequently gave the animals the pellets that were accumulated in the dispenser tube. These sessions were included in the analysis, as the animals received their earned pellets at the end of the session.  (We also checked that excluding the jam sessions did not substantially alter the results or the conclusions.)

### Supplementary Results

| *Control condition* | *Dose of Ex* | *mean difference (kcal)* | *SE (kcal)* | *F or t* | *df* | *p* |
|---|---|---|---|---|---|---|
| Vehicle | | | | 60.21 | 2,8.1 | **$1.3\times10^{-5}$** |
| | 1 µg/kg | −174 | 49 | −3.55 | 4 | 0.057 |
| | 5 µg/kg | −444 | 53 | −8.37 | 4 | **0.0028** |
| | 20 µg/kg | −540 | 20 | −26.58 | 4 | **$3\times10^{-5}$** |
| Baseline | | | | 38.24 | 1.8,7.2 | **0.0002** |
| | 1 µg/kg | −133 | 54 | −2.47 | 4 | 0.16 |
| | 5 µg/kg | −403 | 75 | −5.39 | 4 | **0.014** |
| | 20 µg/kg | −500 | 43 | −11.76 | 4 | **0.0007** |
| Recommended | | | | | | |
| | 1 µg/kg | −145 | 64 | −2.29 | 4 | 0.084 |
| | 5 µg/kg | −415 | 72 | −5.79 | 4 | **0.0044** |
| | 20 µg/kg | −511 | 41 | −12.48 | 4 | **0.0002** |

**Supplementary Table 1. Marginal mean effects of the three doses of Ex on daily total energy intake compared to Vehicle, Baseline and Recommended control in S1 and S2.** First rows for the Vehicle and Baseline control conditions are omnibus F tests, as signified by the two (Greenhouse-Geisser corrected) degrees of freedom. The remaining rows are Dunnett t-contrasts against Vehicle and Baseline, with multiplicity-corrected p-values. The rows under the section labelled ‘Recommended’ are contrasts against the nutritional recommendation with uncorrected p-values.

| *Session* | *Control condition* | *Dose of Ex* | *mean difference (g)* | *SE (g)* | *F or t* | *df* | *p* |
|---|---|---|---|---|---|---|---|
| S1 | Vehicle | | | | 26.36 | 1.6,6.5 | **0.001** |
| | | 1 µg/kg | −57.4 | 10.7 | −5.34 | 4 | **0.015** |
| | | 5 µg/kg | −83.4 | 9.7 | −8.61 | 4 | **0.003** |
| | | 20 µg/kg | −88.2 | 11.3 | −7.83 | 4 | **0.004** |
| | Baseline | | | | 22.82 | 1.8,7.2 | **0.0009** |
| | | 1 µg/kg | −54.6 | 12.3 | −4.44 | 4 | **0.028** |
| | | 5 µg/kg | −80.6 | 10.4 | −7.78 | 4 | **0.004** |
| | | 20 µg/kg | −85.4 | 11.8 | −7.27 | 4 | **0.005** |
| S2 | Vehicle | | | | 22.86 | 1.7,6.8 | **0.001** |
| | | 1 µg/kg | +5.3 | 5.0 | +1.06 | 4 | 0.63 |
| | | 5 µg/kg | −47.5 | 14.3 | −3.32 | 4 | **0.07** |
| | | 20 µg/kg | −70.7 | 9.7 | −7.30 | 4 | **0.005** |
| | Baseline | | | | 20.01 | 1.9,7.7 | **0.001** |
| | | 1 µg/kg | +14.3 | 7.9 | +1.81 | 4 | 0.31 |
| | | 5 µg/kg | −38.5 | 14.4 | −2.68 | 4 | 0.13 |
| | | 20 µg/kg | −61.7 | 8.1 | −7.61 | 4 | **0.004** |

**Supplementary Table 2. Marginal mean effects of the three doses of Ex on food consumption compared to Vehicle and Baseline control in S1 and S2.** First rows for each Session and Control condition combination are omnibus F tests, as signified by the two (Greenhouse-Geisser corrected) degrees of freedom. The remaining rows are Dunnett t-contrasts against Vehicle or Baseline, p values are corrected within each Session and Control condition combination.

**Control analyses of Vehicle and Baseline data**

For the *Dose-finding experiment*, we verified using linear and quadratic contrasts in repeated measures ANOVAs that no significant temporal trends were present in S1 and S2 food consumption across (no-treatment) Baseline control days (Wednesdays of 6 Baseline weeks, S1: p=0.14 and p=0.92, S2: p=0.80 and p=0.40 for linear and quadratic trends, respectively) and across Vehicle control days (3 Tuesdays preceding treatment days, S1: p=0.49 and p=0.26, S2: p=0.67 and p=0.08 for linear and quadratic trends, respectively). We also verified that food consumption in the Vehicle and Baseline control condition did not differ significantly (linear mixed model with random intercepts per subjects, main effect of Control period: $F_{1,39}$=0.12, p=0.73).

For the *Meal schedule experiment*, we compared the Vehicle and Baseline control data to provide further context to the analyses in the main text. Overall food consumption levels did not differ between Vehicle and Baseline either in S1 ($F_{1,4}$=2.33, p=0.20) or S2 ($F_{1,4}$=0.05, p=0.83). The only difference between the two control data was the following: in the Baseline data, S1 food consumption was higher for the more palatable 'very berry' pellets compared to the banana-flavoured pellets (S1 meal type in Baseline: $F_{1,4}$=11.35, p=0.028), while in the Vehicle data there was no significant difference in consumption based on the current pellet type (S1 meal type in Vehicle: $F_{1,4}$=0.29, p=0.62; S1 meal type × Control type interaction: $F_{1,4}$=13.76, p=0.020). However, as the key palatability-based anticipation effect that we focused on was strong in both datasets (S2 meal type main effect across the two datasets: $F_{1,4}$=22.62, p=0.009; see also main text), the above observed subtle difference in the S1 food consumption patterns between the two control conditions does not influence the conclusions of the study.

**Blood glucose level analysis**

We analysed the blood glucose level (BGL) data from samples taken during the *Meal schedule experiment* before the daily drug treatment, before the start of S1 and after S1 (termed as 'pre-treatment', ‘pre-S1’ and ‘post-S1’, respectively). The main text describes the results only briefly. Here in the Supplementary Results, we show the results more comprehensively, with additional statistics supporting our points in the main text.

The factors in the BGL data set were Treatment (Vehicle control, Ex 1 µg/kg), Time (pre-treatment, pre-S1, post-S1), S1 meal type ($b_{S1}$, $v_{S1}$) and S2 meal type ($b_{S2}$, $v_{S2}$). First, we calculated change scores using the pre-treatment value as baseline (pre-S1−pre-treatment, post-S1−pre-treatment), and conducted a repeated measures ANOVA with the above factors, except for Time having only 2 levels (pre-S1 and post-S1). We only report the intercept term from this ANOVA, equivalent to a one-sample t-test of pre-S1 and post-S1 averaged, against baseline. For easier interpretability, we went on to analyse the raw BGL from pre-S1 and post-S1 (again, Time having only 2 levels) obtained in mmol/L (not on change scores) using a mixed model controlling for the (mean-centered) pre-treatment value as a covariate. A linear mixed model (using the `lme4,lmerTest,emmeans` packages in `R`) was fit to the raw BGL data, entering the pre-treatment value as a baseline covariate. Random effect components were chosen so that the model included as many random terms implied by the experimental design as possible, limited by the convergence of the model[52]. Random effects were fit for the intercept, the main effect of S1 meal type, and main effects and all interactions of S2 meal type, Time and Treatment. Meal schedule-related effects (S1 meal type, S2 meal type) were coded as simple contrasts (b=−0.5, v=+0.5), while Time and Treatment effects were coded as dummy contrasts (pre-S1=0, post-S1=1 and Vehicle=0, Ex=1). Thus, the intercept corresponds to the average BGL over all meal schedule conditions in the Vehicle condition’s pre-S1 time point of the Vehicle condition. The contrasts in the results are calculated from this model, unless otherwise noted. Besides the model contrasts, pairwise contrasts and factorial ANOVA-like contrasts are reported (termed ‘sub-models’) to support inferences from the main model contrasts. Degrees of freedom were approximated using Satterthwaite’s method.

In the change-score analysis, we have found that blood glucose levels in general substantially dropped from pre-treatment to pre-S1 ($F_{1,4}$=8.79, p=0.041). The following results are all from the baseline-adjusted mixed model analysis.

Under Vehicle treatment, the pre-S1 BGL was similar across all meal schedules (S1 meal type: $t_{12.1}$=−1.82, p=0.093; S2 meal type: $t_{8.3}$=1.34, p=0.22; S1 meal type × S2 meal type: $t_{50.3}$=1.52, p=0.14). During Vehicle treatment in S1 sessions with very berry flavoured pellets offered, BGL increased (from pre-S1 to post-S1) by 0.76 ± 0.13 mmol/L, a significantly (Time × S1 meal type: $t_{50.0}$=4.30, p=$8\times10^{-5}$) larger increase compared to sessions when banana flavoured pellets were offered, where it stagnated (+0.06 ± 0.13 mmol/L). This effect was erased by Ex treatment (post-S1 minus pre-S1, very berry consumed, Ex treatment: +0.11 ± 0.18 mmol/L), which is not surprising given the similar low consumption rates across conditions under the effect of the drug.

Exenatide had a marked effect on pre-S1 BGL that strongly depended on meal schedules (pre-S1, Treatment: $F_{1,4.2}$=41.50, p=0.0025; Treatment × S1 meal type × S2 meal type: $t_{50.4}$=3.50, p=0.001): the effect was outstandingly strong in the $b_{S1}v_{S2}$ meal schedule condition (d=−2.38 ± 0.25 mmol/L, $t_{6.8}$=−9.7, p=$3\times10^{-5}$, uncorrected contrast) compared to all the other conditions ($b_{S1}b_{S2}$: d=−0.73 ± 0.21 mmol/L, $t_{9.9}$=−3.5, p=0.006; $v_{S1}b_{S2}$: d=−0.24 ± 0.21 mmol/L, $t_{9.6}$=−1.16, p=0.27, $v_{S1}v_{S2}$: d=−0.73 ± 0.25 mmol/L, $t_{6.9}$=−2.97, p=0.021, uncorrected contrasts). In agreement with this, in a sub-model of pre-S1 BGL under Ex treatment, the main effects of S1 meal type ($F_{1,12.1}$=27.1, p=0.0002) and S2 meal type ($F_{1,4.6}$=27.0, p=0.004) and their interaction ($F_{1,50.2}$=42.4, p=$3\times10^{-8}$) were highly significant. These effects substantially weakened and were not significant in the post-S1 BGL samples (sub-model, post-S1 BGL under Ex treatment: S1 meal type: $F_{1,12.1}$=0.04, p=0.84; S2 meal type: $F_{1,4}$=4.98, p=0.077; S1 meal type × S2 meal type: $F_{1,50.2}$=3.39, p=0.071, also supported by the Treatment × Time × S1 meal type × S2 meal type interaction in the main model: $t_{50.0}=-2.61$, $p=0.012$), though note that BGL was still significantly lower under Ex compared to Vehicle control treatment (post-S1, Treatment: $F_{1,4.1}=24.23$, $p=0.0073$).
